# Expression profile of the ADHD risk gene *ADGRL3* during human neurodevelopment and the effects of genetic variation

**DOI:** 10.1101/2025.01.29.635411

**Authors:** Rhiannon Victoria McNeill, Matthias Nieberler, Zora Schickardt, Franziska Radtke, Andreas Chiocchetti, Sarah Kittel-Schneider

**Affiliations:** Department of Psychiatry, Psychosomatics and Psychotherapy, University Hospital Würzburg, Würzburg, Germany; Department of Child and Adolescent Psychiatry, Psychotherapy and Psychosomatics, University Hospital, Würzburg, Würzburg, Germany; Department of Child and Adolescent Psychiatry, Psychosomatics and Psychotherapy, Goethe University Frankfurt, Germany; Department of Psychiatry and Neurobehavioural Science; University College Cork, Cork, Ireland; APC Microbiome, University College Cork, Cork, Ireland

**Author notes:** **Corresponding author:** Rhiannon Victoria McNeill, Department of Psychiatry, Psychosomatics and Psychotherapy, Margarete-Höppel-Platz 1, University Hospital Würzburg, Würzburg 97080, Germany.

## Abstract

Attention-deficit/hyperactivity disorder (ADHD) is one of the most common neurodevelopmental disorders, affecting ∼5.3 % children, with symptoms including hyperactivity, inattention and impulsivity. Moreover, ADHD persists into adulthood in ∼50 % cases, significantly affecting quality of life. Currently, the complex aetiology of ADHD remains unclear, although it has been shown that it is a highly heritable disorder. Single nucleotide polymorphisms (SNPs) in the adhesion G protein-coupled receptor isoform L3 gene (*ADGRL3*/ *LPHN3*) have been consistently associated with ADHD development, symptom severity and treatment response, with the rs1397547 SNP previously shown associated with altered *ADGRL3* transcription. Animal models have revealed an essential role of ADGRL3 in glutamatergic synapse development. However, ADGRL3 function has not been investigated in humans. We used human induced pluripotent stem cell (hiPSC)-derived cortical neurons to characterise ADGRL3 expression during human neurogenesis in vitro, and found that expression peaks early in neurodevelopment, consistent with in vivo data. ADGRL3 protein was found primarily expressed in glutamatergic neurons, and localised to growth cone-like structures, supporting a role in neurite outgrowth and glutamatergic synapse development. Lastly, using hiPSC-derived cortical neurons from healthy controls and ADHD patients, we investigated whether the rs1397547 SNP could affect ADGRL3 expression in cortical neurons. We found the rs1397547 SNP was associated with significantly increased *ADGRL3* transcription in early neurodevelopmental stages. Moreover, single-cell RNA sequencing of maturing cortical neurons revealed a unique transcriptional profile in SNP carriers. Our results further implicate ADGRL3 in ADHD development and suggest that genetic variation may result in dysregulated glutamatergic neuron development.

## Introduction

Single nucleotide polymorphisms (SNPs) in the adhesion G protein-coupled receptor isoform L3 (*ADGRL3*; also known as *latrophilin-3*) gene have been robustly linked to the neurodevelopmental disorder attention-deficit/hyperactivity disorder (ADHD) in both European and non-European populations^1–3^, an association which was further confirmed by a recent meta-analysis^4^. *ADGRL3* SNPs have also been associated with ADHD subtypes, symptom severity and treatment response^1,5^, as well as several other disorders which are frequently comorbid with ADHD such as autism spectrum disorder (ASD), substance use disorder (SUD) and bipolar disorder (BPD)^6^. One of these SNPs is rs1397547 (major allele=C, minor allele=G), which was first identified as associated with ADHD in a well-characterised cohort in 2011^7^. rs1397547 is a synonymous SNP, located in an exon correlating to the seven transmembrane domain. It is classified as a rare genetic variant in European populations due to a minor allele frequency (MAF) of only 0.07, and is located in a highly evolutionarily conserved minimal critical region^8^, suggesting its location is of high biological importance. rs1397547 has not only been associated with ADHD, but was also found associated with an endophenotype characterising ADHD, namely reduced NoGo Anteriorisation value^9^. Additionally, in a previous study we were able to show that rs1397547 genotype was associated with significantly increased *ADGRL3* transcription in human fibroblast cells^10^, further highlighting its biological importance.

The physiological role of ADGRL3 remains largely unknown, particularly in humans. Animal models have revealed that ADGRL3 is an adhesion G protein-coupled receptor (aGPCR) thought to be involved in regulating intracellular calcium concentration^11^, neurite outgrowth^12^, and the development of glutamatergic synapses^13^. It is a constitutively active aGPCR that increases intracellular cAMP levels, and this activity is thought to be essential for promoting synapse formation^14^. It is currently unclear whether it is a pre-synaptic or post-synaptic receptor, but potentially both, depending on developmental stage and cell type^6^. Mouse Adgrl3 has been observed to bind to Unc5D^15^, Ten1^16^ and Flrt3^13^, which together form a combinatorial code to cause adhesion or repulsion during neuronal migration and mediate synapse specificity^17^. Interestingly, CNVs in the human *FLRT3* gene have also been associated with ADHD^18^. ADGRL3 knock out (KO) models in mice showed consequent hyperactivity^19,20^, calcium signalling changes^12^, and a 40% loss of glutamatergic synapses^21^, with similar phenotypes found in KO rats^22^. In zebrafish, ADGRL3 KO also resulted in a hyperactive phenotype, which could be rescued by methylphenidate treatment^23^. While these KO models have been essential for identifying ADGRL3 function, they unfortunately lack construct validity, as none of the identified *ADGRL3* genetic variants implicated in ADHD cause a complete gene deletion^24^.

Expression of ADGRL3 in humans has currently only been investigated using post-mortem brain tissue, with particularly high expression observed in the amygdala, cerebellum and cerebral cortex, all of which are regions implicated in ADHD^1^. ADGRL3 differs to other isoforms (ADGRL1 and ADGRL2) as it is the isoform which accounts for most of the genetic variants found associated with human neuropsychiatric disorders^6^. However, its functional role currently remains unclear, with very few studies conducted to investigate ADHD-related *ADGRL3* genetic variations at the molecular level, particularly in humans. Elucidating the role of ADGRL3 and the effects of genetic variation are therefore critical to understanding ADGRL3’s potential pathological function in neuropsychiatric disorders, particularly ADHD.

In this study, we performed a preliminary investigation into the expression prolife of ADGRL3 during human neurodevelopment, and potential effects of the ADHD risk SNP rs1397547. To do this we used human induced pluripotent stem cells (hiPSCs) derived from healthy controls and adult ADHD patients which had different rs1397547 genotypes^25^. hiPSCs maintain the unique complex genotype of the donor they are derived from, are human-based, and provide the opportunity to investigate risk genes in the cell type in which they are thought to exert their specific effects, making them highly suitable for the study of neuropsychiatric disorders^26^. We differentiated hiPSCs into neuronal progenitor cells (NPCs) and cortical neurons (CNs), as ADGRL3 had been reported as highly expressed in the cerebral cortex in human post-mortem brain tissue^1^, and important for glutamatergic synapse development in mice models^21^. Moreover, accumulating evidence implicates an important role of the cortex in ADHD development^27^. We then investigated ADGRL3 gene and protein expression using various methods, including single-cell RNA sequencing (scRNA-seq), to establish a human-specific expression profile and assess the effects of genetic variation associated with ADHD development.

## Materials and Methods

### Participant Recruitment

Healthy participants were primarily employees or employee relatives, and had no previous or current psychiatric, internal, infectious, or severe neurological disorders. They were screened using the Mini-DIPS questionnaire to exclude the presence of a psychiatric disorder. Adult ADHD patients were assessed using the DSM-IV or DSM-5 diagnostic criteria by two independent psychiatrists. Patients were recruited in 2011-2018 at the Department of Psychiatry, Psychosomatics and Psychotherapy, University Hospital Würzburg (Germany) and the Department of Psychiatry, Psychosomatic Medicine and Psychotherapy, University Hospital Frankfurt (Germany). Ethical approval was obtained from the ethics committees at the University of Würzburg (#96/11) and University of Frankfurt (#425/14). Donor demographic data is given in Supplementary Table 1.

### Genotyping

Genomic DNA was extracted from blood samples, fibroblast cells and hiPSCs from donors using the DNeasy kit (Qiagen). A Competitive Allele-Specific PCR (KASP™) assay (LGC) was used to determine rs1397547 genotype in the *ADGRL3* gene.

### hiPSC Generation and Maintenance

hiPSCs were generated and maintained as previously described^28,25^. Briefly, fibroblasts cells were reprogrammed to hiPSCs using the CytoTune-IPs 2.0 Sendai Reprogramming Kit (Invitrogen) and quality controlled (QC). QC included assessment of pluripotency, genetic integrity, confirmation of cell line identification, and absence of viral genome and mycoplasma. hiPSCs were media changed every other day using StemMACS™iPS-BrewXF (Miltenyi Biotech) and grown on Matrigel™ (Corning)-coated cultureware (diluted according to manufacturer’s instructions). Cells were split when approaching approximately 80% confluency using RELESR™ (Stemcell) in a 1:6 ratio. hiPSCs were cultured in an incubator at 37 °C, 5 % CO_2_ and 20 % O_2_. A minimum of n=3 donors was used for experiments, with the exact sample size for each experiment given in figure legends.

### Neural Progenitor Cell (NPC) Generation and Maintenance

hiPSCs were differentiated into NPCs are previously described^29^. Briefly, hiPSCs were differentiated using dual SMAD inhibition. NPCs were grown on Matrigel diluted 1:40, maintained in Neural Expansion media (Merck) and passaged using Accutase (Stemcell) every 3-5 days at 1:3 ratio.

### Cortical Neuron (CN) Differentiation

NPCs were differentiated into CNs as previously described^29^. Briefly, NPCs were seeded on poly-L-ornithine/laminin (10 µg/mL) at a density of 30 000 cells/cm^2^ and maintained in Neuronal Differentiation media (Merck). For the first two weeks, an 80% media change was performed 3x/week. After this, a 50% media change was performed 2x/week. This protocol results in a mixed cortical neuronal culture after 14 days, consisting predominantly of glutamatergic neurons (∼90%)^29^.

### RT-qPCR

RNA was extracted from cells using the RNeasy-Plus Mini kit (Qiagen), checked for genomic DNA contamination and cDNA synthesis performed using the iScript cDNA Synthesis Kit (Bio-Rad). For amplification, Taqman primers (Thermofisher) were used (see Supplementary Table 2). Amplification was performed on a BioRad CFX machine (BioRad). Data was normalised to two reference genes based on stability plots, and relative transcription of target genes was calculated (^ΔΔ^Ct).

### Immunofluorescent (IF) Labelling

Cells were fixed in 4 % paraformaldehyde (Roth), blocked using 5 % FBS+1 % BSA, and permeabilised using 0.2 % Triton X-100 (Sigma). For labelling of ADGRL3, Triton X-100 was omitted. Primary antibodies were diluted and incubated on samples overnight at 4 °C. Samples were washed 3x with PBS and incubated with secondary antibodies for 1 hour at room temperature. Samples were again washed before staining with DAPI. For details of antibodies see Supplementary Table 3. IF widefield images were captured using the Eclipse Ti2-E epifluorescence microscope (Nikon), confocal images were captured using the AX-R confocal microscope (Nikon). Imaging analysis was performed using the NIS-Elements software (Nikon), and a minimum of 10 images per donor from different XY locations were analysed for each experiment. For widefield images, the Clarify.ai function of NIS-Elements was used to exclude cells on a different focal plane.

### Western Blot

Cells were washed 1x with cold PBS, incubated briefly in N-PER Extraction Reagent (ThermoFisher) supplemented with Protease Inhibitor Cocktail (Roche) and PhosStop (Roche), and scrape-harvested into microtubes. Samples were incubated for 30 minutes on ice with occasional vortexing. Microtubes were centrifuged at 12 000 rpm for 30 minutes at 4 °C, and supernatant transferred to a new microtube. Protein concentration was measured using the ADV02 assay (Cytoskeleton Inc) and a Tecan SPARK plate reader (Tecan). 20 μg protein was separated using a 4-12 % Bolt™ Bis-Tris gel in Bolt™ MOPs buffer (Thermofisher), transferred onto a Nitrocellulose membrane (pore size 0.45 μM; ThermoFisher) and membranes incubated in Intercept® blocking buffer (LI-COR) for 1 hour at room temperature. Primary antibodies were diluted in a PBST/blocking buffer solution and incubated on membranes overnight at 4 °C on a rocker. Membranes were then washed with PBST on a rocker, for 3 x 5 minutes. Secondary antibodies were diluted and added to the membranes for 1 hour at room temperature. Membranes were washed and imaged using a Fusion FX imager (Vilber). Total protein was stained using Revert™ total protein stain (LI-COR) and protein quantification performed using Image Studio™ Lite v5.0 software (LI-COR).

### scRNA-seq

8-week-old hiPSC-derived CNs were washed with PBS and dissociated into single cells by incubation with accutase at 37°C for 20 minutes. Remaining cell aggregates were removed using a 30 μM pre-separation filter (Miltenyi). Cells were blocked and then labelled with TotalSeqA Hashtag antibodies (BioLegend) to enable multiplexing. After washing, live cell concentrations were adjusted to 1500 cells/μL. Cells from different donors were then pooled in equal parts and libraries generated using 3’GEX v3.1 (10x Genomics). Sequencing was performed with the NextSeq 2000 platfom (100 cycles, P3 flow-cell; Illumina) and data demultiplexed using the Cell Ranger software (10x Genomics). All subsequent analyses were performed in R v. 4.2.2 using the Seurat Package. For details please see Supplementary Information.

### Data Mining from Public Biorepositories

RNA sequencing data from the BrainSpan developmental transcriptome (accessed: 01/05/2024) was used to determine in which brain region and at what time points *ADGRL3* transcript was expressed during human neurodevelopment, and to identify co-expressed genes^30^. We also used the STAB repository (Spatio-Temporal Cell Atlas of the Human Brain; accessed: 02/05/2024) to further investigate the neurodevelopmental timing of *ADGRL3* expression, and to identify cell subtype-specific expression^31^.

### Statistical Analyses

Statistical analyses were performed using Graphpad Prism 10 (Graphpad). The mean of n=≥2 technical replicates was calculated for each donor. All data were first checked for normal distribution, and parametric or non-parametric tests used accordingly. The specific statistical test performed for each experiment is given in figure legends. The level of significance was set at *p*<0.05. hiPSC lines from different donors are represented as individual data points in figures.

## Results

### ADGRL3 Expression during Human Neurodevelopment

We first investigated the expression profile of the *ADGRL3* gene during human neurodevelopment, utilising in vivo public data repositories and our own hiPSC-based in vitro model. Using BrainSpan, *ADGRL3* transcription was found primarily localised to the dorsolateral prefrontal cortex (DLPFC; Figure 1A). Three primary peaks in transcription were found, all during prenatal development, corresponding to post-conception weeks (PCW) 12, 16 and 24. After birth, *ADGRL3* transcription was observed to decrease. The top co-expressed genes were identified as *NAV3* (*neuron navigator 3*), *NCOA1* (*nuclear receptor coactivator 1*), *SEMA6D* (*semaphorin 6D*), and *SLC8A3* (*sodium-calcium exchanger 3*). scRNA-seq data from the STAB atlas supported this observation, with *ADGRL3* transcript in the DLPFC found to substantially increase prenatally (P5-P6, corresponding to 16-24 PCW) before levelling off after birth (P12; Figure 1B). During this developmental period, it was found that the DLPFC consists of predominantly of excitatory neurons (ExN1_4; Figure 1C), suggesting that the *ADGRL3* gene expression may potentially be localised to this specific cell type. We next investigated the expression profile of ADGRL3 in our hiPSC-derived cortical neuron model, which reflects early embryonic human neurodevelopment^29^, to determine whether this would reflect the in vivo data. We observed a significant increase in *ADGRL3* transcription during differentiation from hiPSCs into NPCs, with transcription peaking in 2-week-old CNs (days in vitro 14/DIV14) and remaining high in 8-week-old cultures (DIV56; Figure 1D). ADGRL3 protein expression supported this trajectory, with peak expression observed in NPCs and 2-week-old CNs (DIV14), before significantly decreasing in 8-week-old cultures (DIV56) cultures (Figure 1E). This suggested that our in vitro model was able to reflect the early prenatal expression profile of *ADGRL3* found in vivo, and was suitable for further characterisation of ADGRL3 expression.

**Figure 1.**
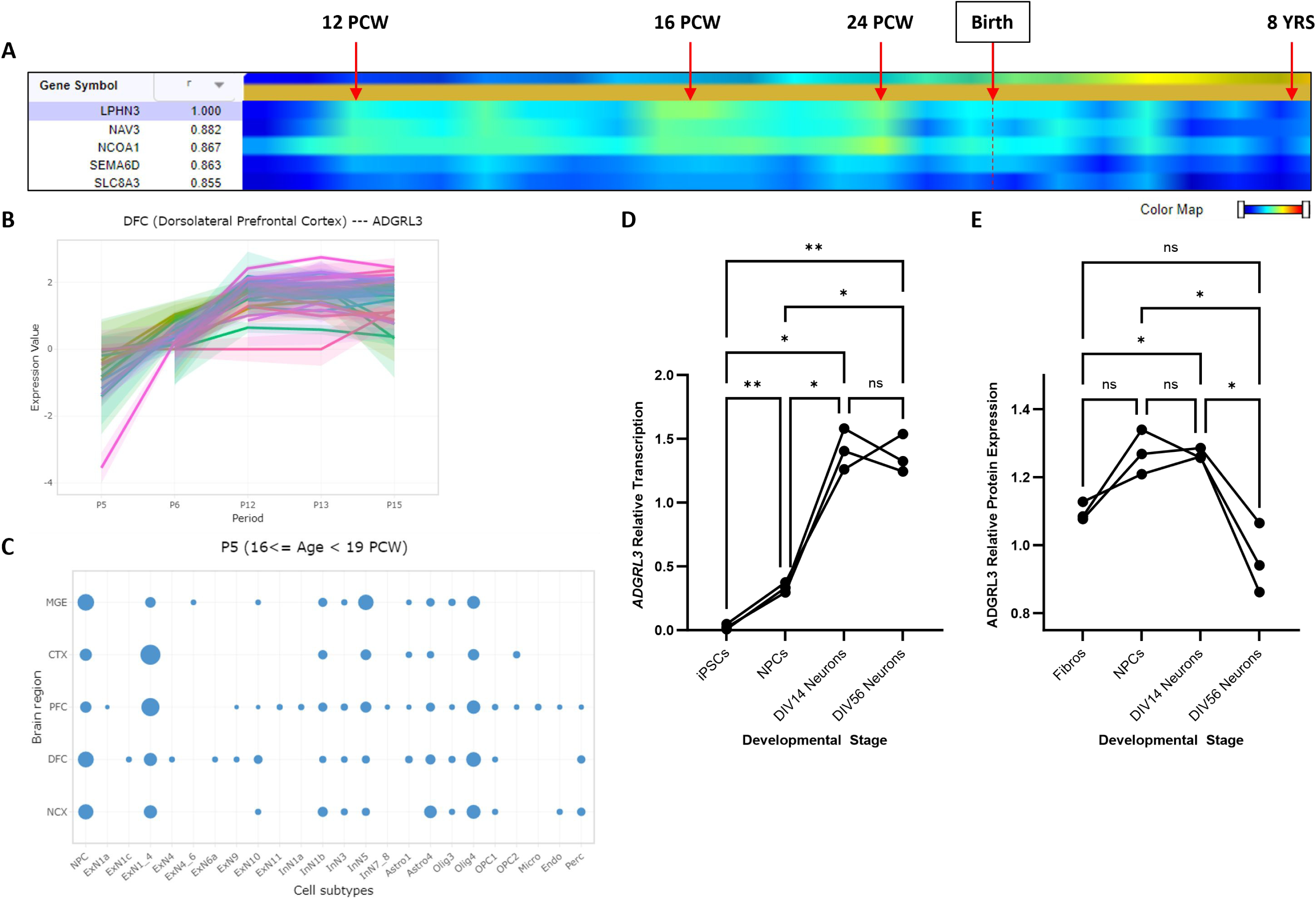
ADGRL3 expression during human neurodevelopment. (A) Heatmap showing *ADGRL3* transcript expression in the DLPFC at pre- and post-gestational time points using the the BrainSpan developmental transcriptome atlas. Expression data for the top 5 co-expressed genes are also shown, along with the correlation coefficient (r). B) *ADGRL3* transcript expression in the DLPFC at different periods (P) post-conception, extracted from the STAB database. P5: 16<= Age < 19 PCW; P6: 19<= Age < 24 PCW; P7: 24<= Age < 38 PCW; P12: 12<= Age < 20 Years; P13: 20<= Age < 40 Years; P14: 40<= Age < 60 Years; P15: > 60 Years. (C) Cell type composition of the developing brain at P5. Larger circle = higher proportion of cell type. (D) *ADGRL3* transcription during hiPSC-derived neuronal differentiation, assessed using RT-qPCR (n=3). (E) ADGRL3 protein expression during hiPSC-derived neuronal differentiation, quantified using western blot (n=3). Significance was tested using repeated measures one-way ANOVA. PCW=post-conceptional week; MGE=medial ganglionic eminence; CTX=cortex; PFC=prefrontal cortex; DFC=dorsolateral prefrontal cortex; NCX=neocortex; hiPSCs=human induced pluripotent stem cells; NPCs=neural progenitor cells; DIV=days in vitro.

### ADGRL3 Receptor Localisation

We next set out to determine the localisation of ADGRL3 protein, which could help infer its role in human neurodevelopment. As we observed peak ADGRL3 expression in 2-week-old CNs, we used these cells to investigate ADGRL3 localisation using IF and co-labelling with neuronal markers. Firstly, we identified that ADGRL3 was colocalised to TUBB3-positive (TUBB3+) axons, confirming its expression in neurons (Figure 2A). ADGRL3 protein was found located both along axonal segments, but also appeared highly expressed at axon ends in growth cone-like structures. We additionally observed ADGRL3 expression in cells weakly positive for TUBB3, specifically in microtubular structures (Figure 1B). We next investigated whether ADGRL3 could be found specifically localised in excitatory neurons as suggested by the in vivo data, using the glutamatergic neuron marker vGLUT2. IF images revealed that ADGRL3 localised to vGLUT2+ axons, along axonal segments and again in growth cone-like structures (Figure 2C). We also observed ADGRL3+ axons which did not contain vGLUT2, and vice versa (Figure 2D).

**Figure 2.**
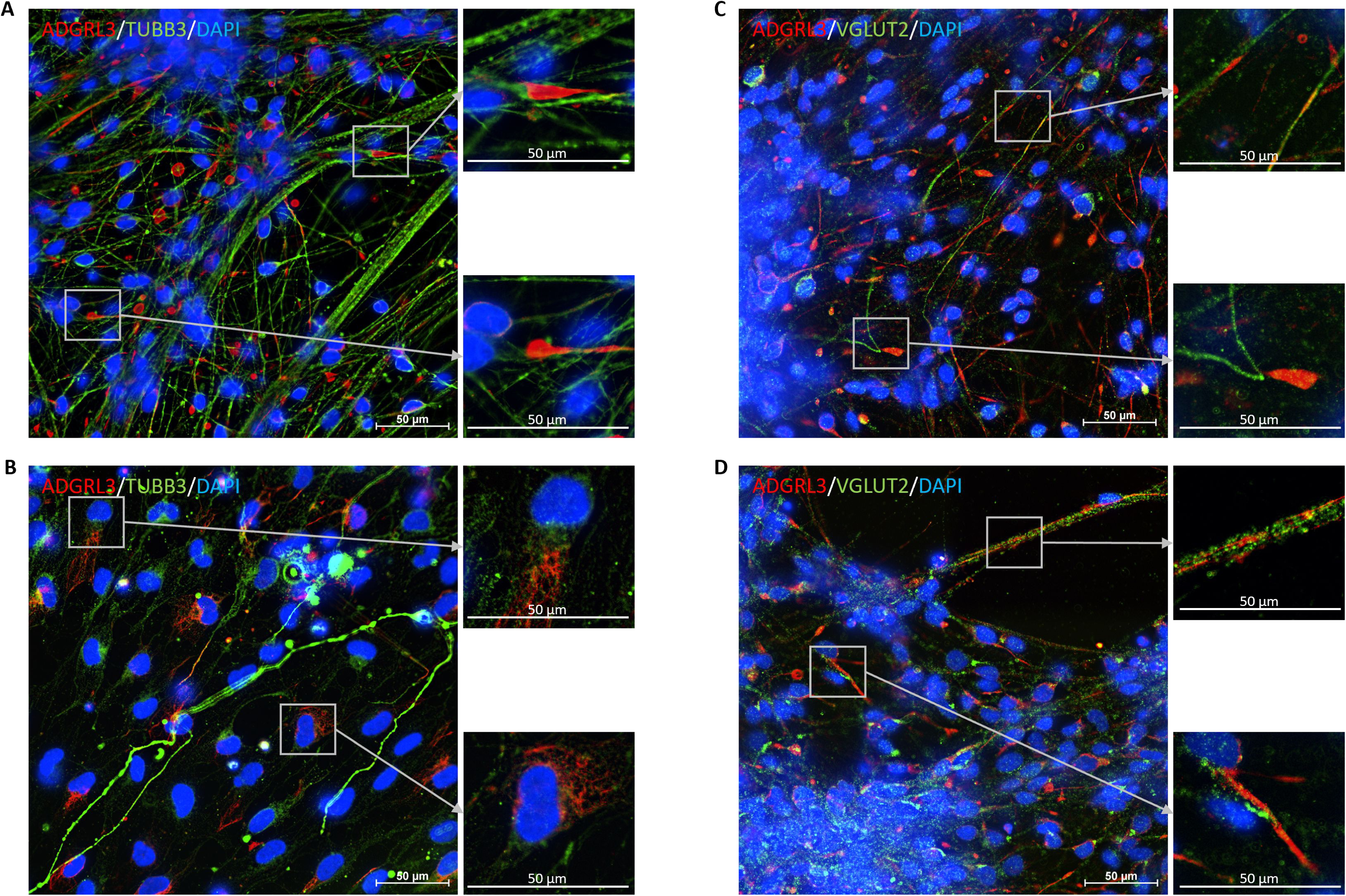
ADGRL3 localisation in hiPSC-derived CNs. hiPSC-derived NPCs were differentiated into CNs for 2 weeks (n=12). Immunofluorescence (IF) labelling was then performed for ADGRL3, the neuron-specific marker TUBB3 and the glutamatergic neuron marker vGLUT2. Representative images are shown. (A+B) ADGRL3 was found expressed in TUBB3+ cells. Specifically, ADGRL3 was found (A) highly expressed at the terminal end of TUBB3+ axons in growth cone-like structures, and (B) within microtubule-like structures. (C) ADGRL3 was found expressed in vGLUT2+ axons, appearing to occasionally colocalise. (D) ADGRL3+ axons were also observed that did not express vGLUT2.

To further investigate the localisation of ADGRL3, we also performed confocal microscopy and 3D rendering of 2-week-old CNs from wild type healthy controls. IF labelling for TUBB3 and ADGRL3 further confirmed the axonal localisation of ADGRL3, but also revealed expression on the surface of nuclei (Figure 3A). We could also observe the presence of TUBB3+ axons which did not appear to contain ADGRL3. 3D rendering supported earlier 2D results of intense ADGRL3 labelling in growth cone-like structures, as shown in the magnified image (denoted by arrows). Interestingly, we were also able to observe a distinct apical-basal patterning of ADGRL3 expression, whereby ADGRL3 was highly basally distributed (Figure 3B). IF labelling for ADGRL3 and vGLUT2 also confirmed the 2D results, whereby ADGRL3+ axons were also co-labelled for vGLUT2 (Figure 3C). The observed results from 3D rendering were also replicated in a second cell line used, particularly the basal distribution of ADGRL3 expression (Supplementary Figure 1).

**Figure 3.**
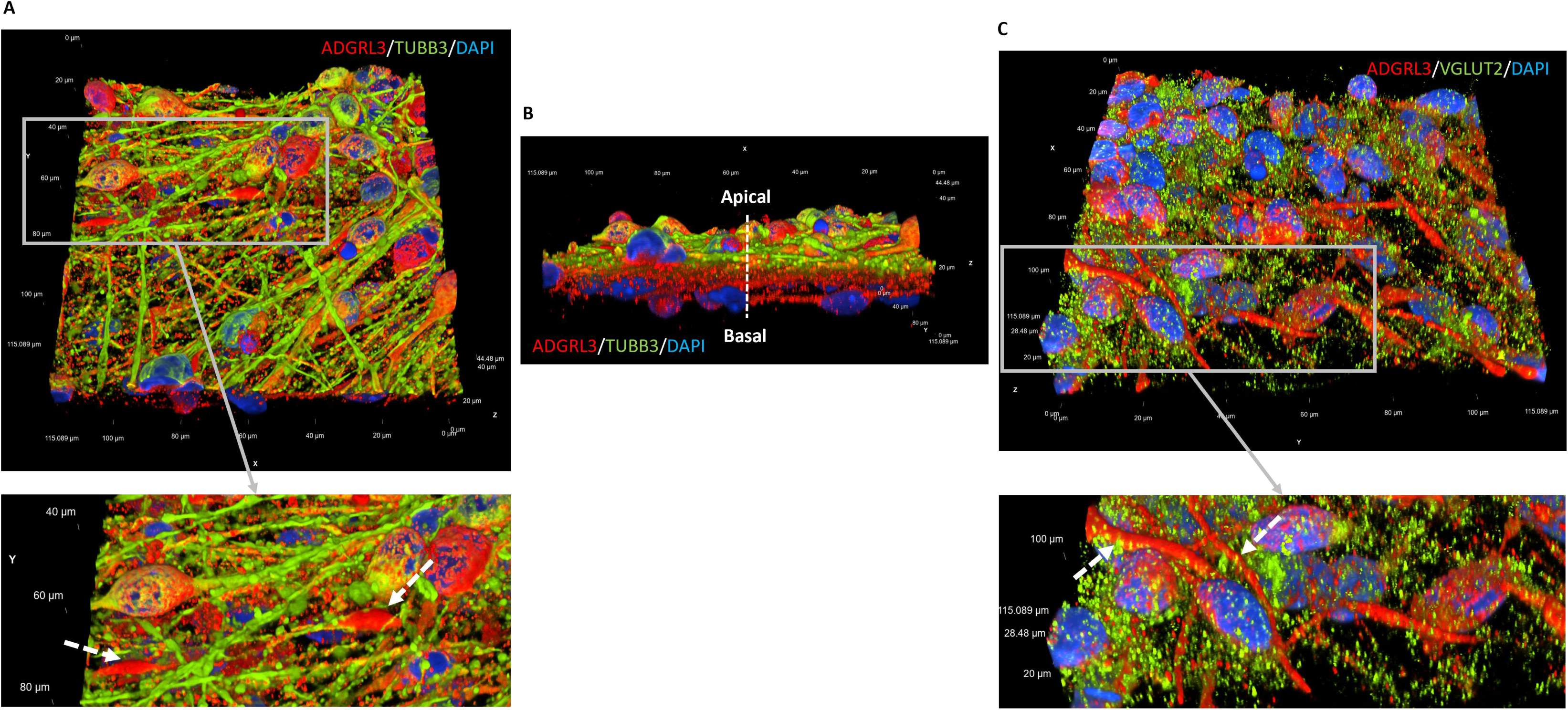
3D localisation of ADGRL3 in hiPSC-derived CNs. IF was carried out on 2-week-old CNs and cells were imaged using confocal microscopy (n=2). 3D rendering was then performed. (A) Representative images for ADGRL3 and TUBB3 labelling, with a region of interest containing growth cone-like structures further magnified (denoted by arrows). (B) 3D rendering showed a distinct basal patterning of ADGRL3 expression. (C) Representative images for ADGRL3 and vGLUT2 labelling. A region of interest containing ADGRL3+ axons colocalised with vGLUT2 is magnified (denoted by arrows).

### Effects of rs1397547 on Gene Expression

We next investigated whether the ADHD risk SNP rs1397547 could affect ADGRL3 expression. As ADGRL3 expression was found to rapidly increase during differentiation into NPCs and 2-week-old CNs, we focused on these developmental stages. *ADGRL3* transcription was found significantly increased in both NPCs (Figure 4A) and two-week-old CNs (Figure 4B) from donors carrying at least one copy of the rs197547 SNP. However, no significant changes were observed in protein expression (Figure 4C&4D). To exclude the possibility that the observed increase *ADGRL3* transcription was due to a higher number of neurons expressing *ADGRL3*, we quantified the percentage of TUBB3 and vGLUT2-positive areas in CNs also expressing ADGRL3, as well as ADGRL3 fluorescence intensity, and found no changes between genotypes (Supplementary Figure 2A&2B).

**Figure 4.**
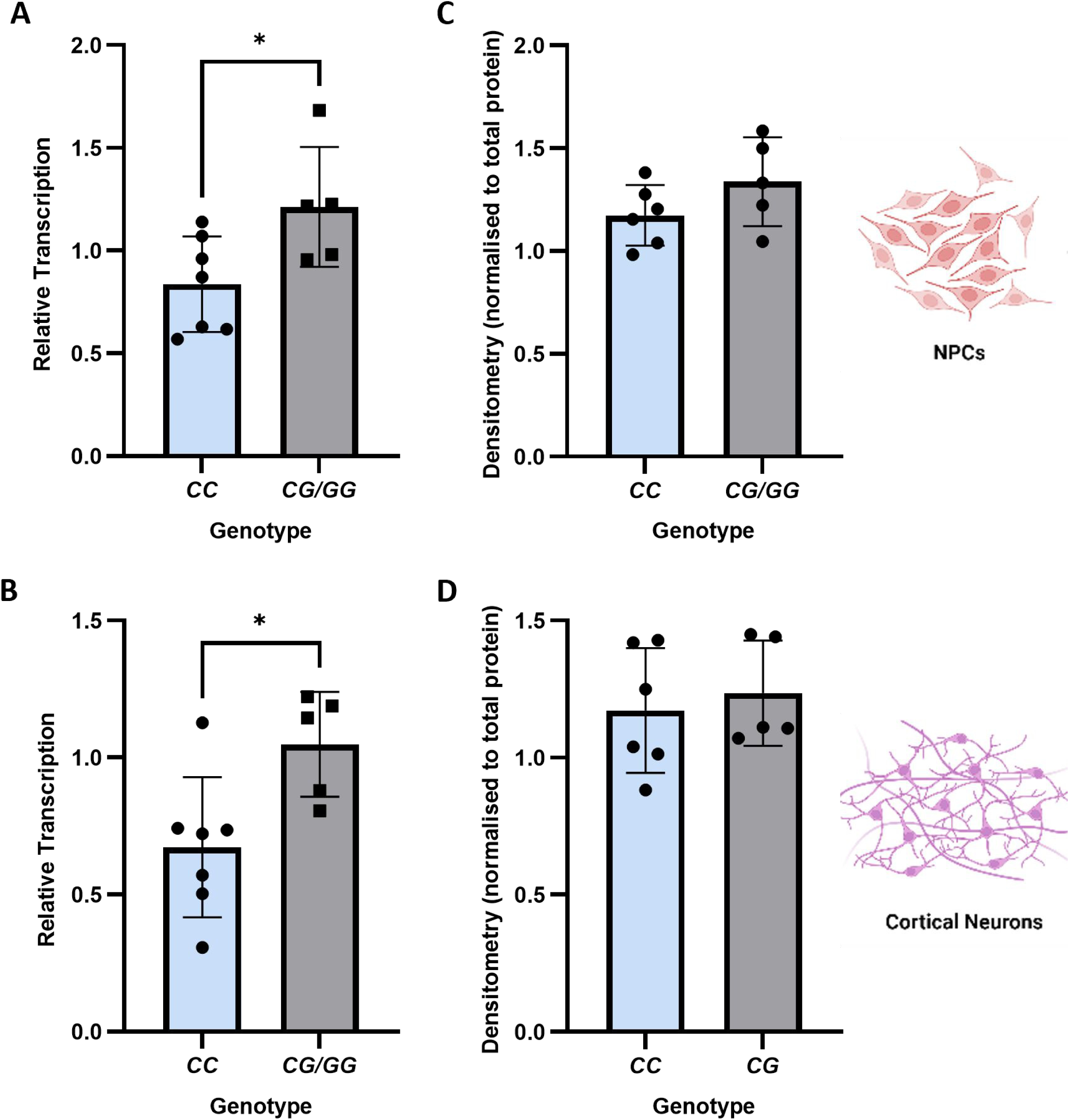
The rs1397547 SNP is associated with altered ADGRL3 expression. RT-qPCR was performed to quantify relative transcription, and western blotting to quantity protein expression, in both (A) NPCs and (B) 2-week-old CNs (n=7 CC, n=5 CG/GG). Significance was determined using unpaired t-tests. NPCs=neural progenitor cells.

We additionally investigated later developmental time points (4- and 12-week-old CNs), but rs1397547 genotype-associated changes in *ADGRL3* transcription were no longer observable (Supplementary Figure 3). Despite this, genotype-associated changes in the expression of other glutamatergic synapse-related genes could be observed in functionally mature 12-week-old CNs. Data were split according to diagnosis and rs1397547 genotype, to exclude the possibility that the changes observed were solely due to ADHD diagnosis. The results revealed a significant downregulation of *vGLUT1* transcription in ADHD patient rs1397547 carriers compared to ADHD wild type (Figure 5A), and a linear decrease in *PSD95* transcription depending on diagnosis and rs1397547 genotype (Figure 5B). Lastly, a significant upregulation of *EAAT1* transcription was found in rs1397547 carriers compared to both wild type healthy controls and wild type ADHD patients (Figure 5C).

**Figure 5.**
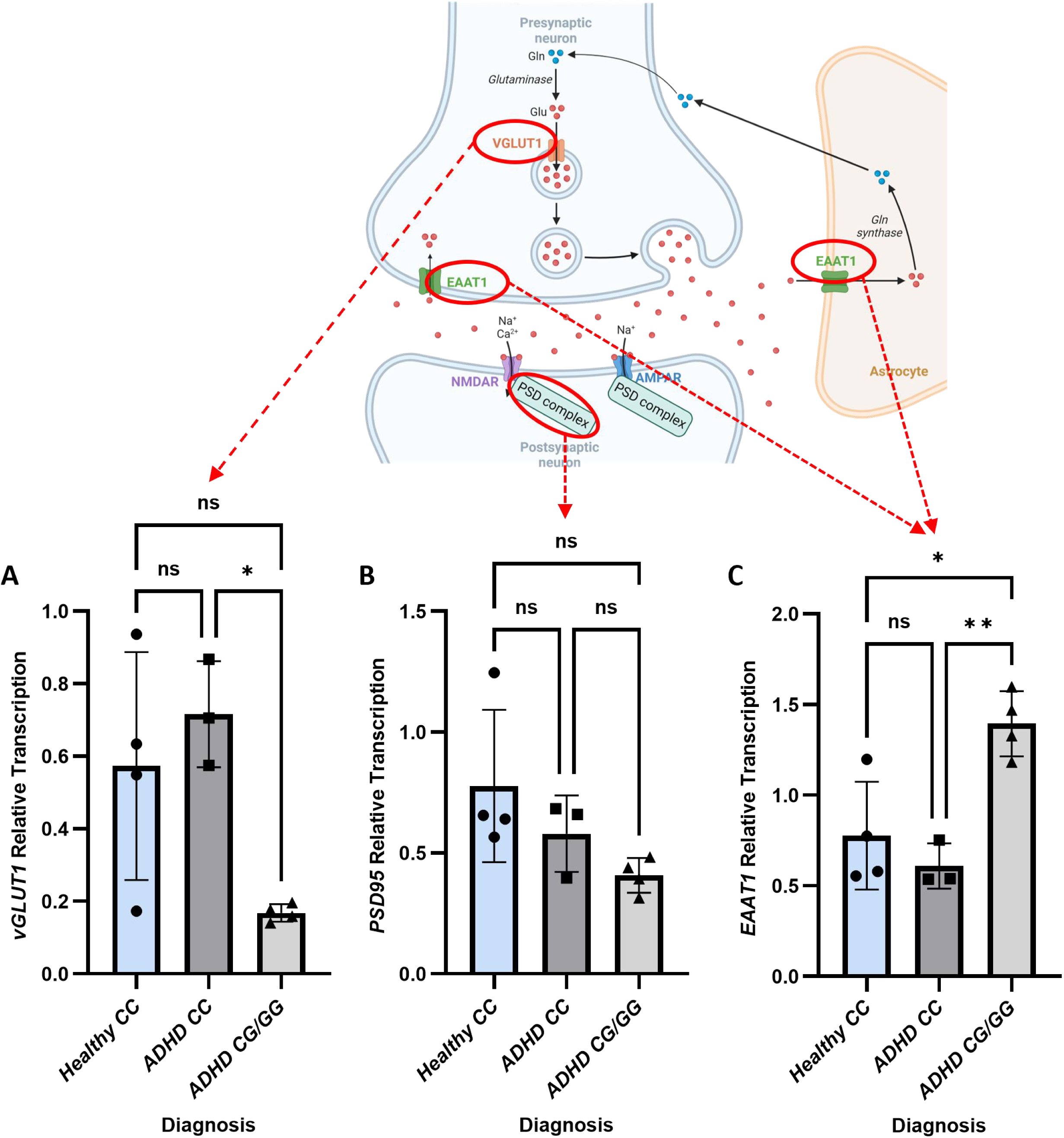
Effects of the rs1397547 SNP on glutamatergic synapse-related gene expression in functionally mature neurons. CNs were differentiated for 12 weeks, and RT-qPCR performed. Data were split according to diagnosis and genotype (n=4 healthy control CC, n=3 ADHD CC, n=4 ADHD CG/GG). Relative transcription was calculated for (A) *vGLUT1*, (B) *PSD95* and (C) *EAAT1*. Significance was tested using one-way ANOVA.

### Cell-specific *ADGRL3* gene expression and effects of genetic variation using scRNA-seq

scRNA-seq data from 8-week-old cortical neurons was generated as part of a different project, however we utilised the available data to provide further insights into ADGRL3 in human neurodevelopment. Clustering analysis revealed a total of 19 different cell types present in our cortical neuron cultures, the majority of which were cortical neural lineages (Supplementary Figure 4A). *ADGRL3* was found to be most highly expressed in clusters 7 and 13 (Figure 6A), along with *MAP2* and *GRIN2B*, which were identified as layer 2/3 neurons (L23) and cycling ventricular radial glia (ccvRG) respectively (Supplementary Figure 4B). Gene ontology analysis showed that cluster 7 was significantly associated with synaptic membrane, neuron projection and trans-synaptic signalling, whereas cluster 13 was associated with mitotic cell cycle (Figure 6B). Cell type distribution analysed showed that rs1397547 SNP carriers had significantly less cluster 7 and 13 cells compared to wild type (Figure 6C).

**Figure 6.**
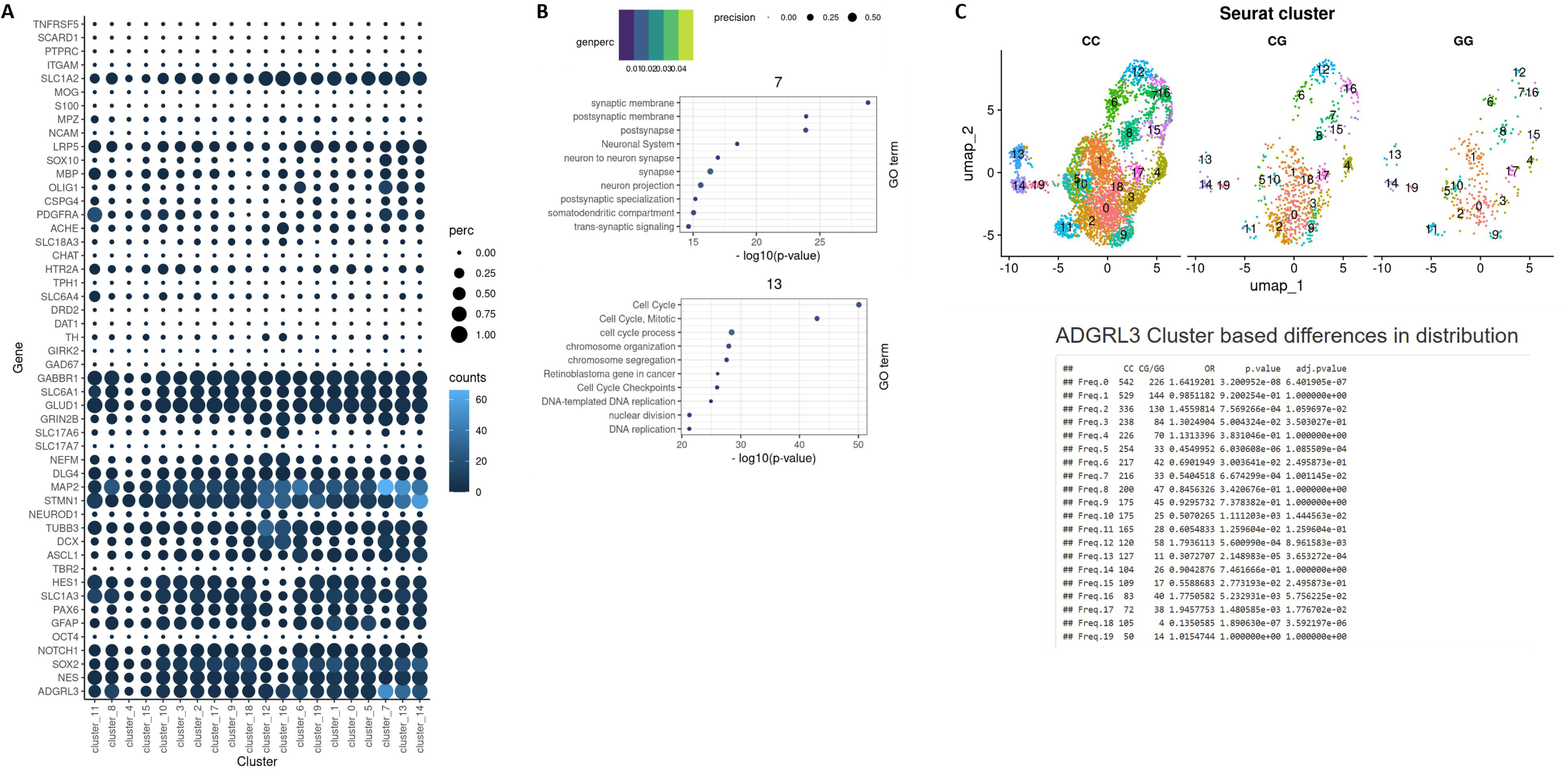
Expression of *ADGRL3* in specific neuronal subtypes and transcriptomic changes associated with rs1397547 genotype. scRNA-seq was performed using 8-week-old CNs (n=4 HC, n=5 ADHD; n=6 CC, n=3 CG/GG). UMAP clustering was performed and cell types identified using brain development reference data. (A) Expression of cell type-associated genes in the different clusters, with *ADGRL3* most highly expressed in clusters 7 and 13. (B) Gene ontology analysis for cluster 7 and 3. (C) Cell type distribution was significantly different between rs1397547 genotypes for clusters 7 (adjusted p value=0.01; OR=0.54) and 13 (adjusted p value=0.0004; OR=0.3).

Pseudo bulk gene expression analysis across all cell types revealed several differentially expressed genes between genotypes (Supplementary Table 4), including *MBP* (adjusted p value=1.81×10^-33^; fold change=-2.12), *DSCAML1* (adjusted p value=1.22×10^-22^; fold change - 1.22) and *SLC8A3* (adjusted p value=5.6×10^-15^; OR=-2.22). Cell type-specific DEG analysis identified several genes as differentially expressed due to genotype in cluster 7 (Supplementary Table 5), including *THBS1* (adjusted p value=1.27×10^-7^; fold change=1.85), *GPR37* (adjusted p value=0.000949; fold change=7.64), *IGF*-1 (adjusted p value=0.008713; fold change=2.7) and *MARCKS* (adjusted p value=0.033289; fold change=-1.38). No significant differentially expressed genes were found in cluster 13.

## Discussion

Our study aimed to provide a preliminary investigation into the human-specific expression of the ADHD risk gene *ADGRL3* using hiPSC-derived CNs, along with the potential effects of a rare SNP associated with several ADHD clinical parameters. Comparing our in vitro data with in vivo data available from public biorepositories, we were able to show that our model can recapitulate the developmental timing of *ADGRL3* expression, which shows a significant increase in transcription in early and mid-gestation in the DLPFC. This is consistent with its proposed role in neurite outgrowth^12^ and synapse development^13^, as observed in rodent models. Analysis of top co-expressed genes identified other neuronal guidance proteins, such *NAV3*^32^, in which genetic variants have recently been associated with neurodevelopment disorders^33^. Interestingly, *SEMA6D* was also identified as highly co-expressed, which is a protein important for neuronal wiring and involved in growth cone collapse^34^. *SEMA6D* was also one of the risk loci found significantly associated with ADHD in the first genome-wide association study (GWAS)^35^, and educational attainment in a different GWAS^36^. Lastly, the sodium/calcium exchanger 3 (*SLC8A3*) gene was also a co-expression partner, and is involved in maintaining intracellular calcium homeostasis^37^, further supporting the proposed role of ADGRL3 in calcium signalling^11^.

We next went on to investigate the localisation of ADGRL3 using 2-week-old CNs, as expression was found to most rapidly increase at this time point. Using IF, we were able to confirm that ADGRL3 located to TUBB3+ axons, and that ADGRL3+ axons often contained vGLUT2, supporting previous observations in mice that ADGRL3 is predominantly expressed in glutamatergic neurons^13^. We also observed that TUBB3, vGLUT2 and ADGRL3 did not always colocalise, suggesting a potential temporal order of expression. ADGRL3 protein was often not distributed equally along the length of axons, instead appearing in individual axonal segments, suggesting a specific spatial distribution. The most intense ADGRL3 labelling was observed in growth cone-like structures. This is supported by previous findings of an enrichment of Adgrl3 in growth cones in mice, accompanied by actin remodelling proteins and proteins from neurotransmitter vesicle release machinery such as Snap25^6^. An active role of Adgrl3 in regulating the formation of filopedia and lamellipodia was also found in this study. A more recent paper found ADHD-associated missense mutations impaired intrinsic G13-based aGPCR signalling, which is associated with actin cytoskeleton remodelling, and reported decreased cellular area and perimeter, as well as decreased number of filopedia and lamellipodia^24^. The authors concluded that *ADGRL3* variants interrupt GPCR G13 signalling, resulting in deficient actin cytoskeleton remodelling which may inhibit age-appropriate neurodevelopmental features. We also found ADGRL3 localised in microtubule-like structures, further supporting a potential role for ADGRL3 in cytoskeleton remodelling.

3D rendering of confocal images additionally revealed a distinct spatial pattern of ADGRL3 expression not previously reported. We observed that there appeared to be an apical-basal expression of TUBB3 and ADGRL3, with ADGRL3 expression basally distributed. This observation was made in both cell lines investigated, suggesting it was not due to a technical artefact. This finding could imply a direct interaction of ADGRL3 with the extracellular matrix protein laminin, on which the neurons were cultured, to regulate cell adhesion. Alternatively, laminin could be acting as an extracellular signalling cue to indirectly cause a distinct spatial pattern of ADGRL3 distribution. It has been proposed that laminin plays a role in axonal growth at the basal surface of the neuroepithelium^38^, and that the basal neuroepithelium is a major site of neurogenesis for the neocortex^39^. Given ADGRL3’s proposed role in neurite outgrowth and axon guidance, these factors could account for its basal expression.

We next investigated the molecular consequences of the rs1397547 SNP, focusing on gene expression. At the developmental time points where *ADGRL3* transcription was most upregulated, we found that genotype altered *ADGRL3* transcription, with significantly increased transcription found in donors carrying at least one copy of the rs1397547 SNP. This is consistent with our previous findings in human fibroblast cells^10^. However, we did not find any correlating changes in ADGRL3 protein expression via western blot or IF. This could be because both methods are only semi-quantitative, and western blot in particular may lack the sensitivity to detect modest changes in expression^40^, especially given that protein levels tend to be more conserved than transcript levels^41^. It has also been reported that there is a delay in translation of transcript level changes to protein, particularly during differentiation^42^, which could mean that changes in ADGRL3 protein expression may not be detectable until a later time point. Lastly, it is also possible that post-transcriptional regulation intercedes to maintain homeostatic protein expression of ADGRL3^43^.

To determine whether the changes identified in *ADGRL3* transcription associated with rs1397547 genotype were also present in functionally mature CNs, we investigated gene expression in 12-week-old CNs. Interestingly, genotype-related changes in *ADGRL3* transcription were no longer detectable at this time point, highlighting the importance of studying risk gene expression not only in relevant cell types, but also at relevant developmental time points. However, alterations in the expression of glutamatergic synapse-related genes were found. To exclude the possibility that these changes were due to ADHD diagnosis, we separated the data based on diagnosis and rs1397547 genotype. This revealed a unique transcriptional signature specific to ADHD patients carrying the rs1397547 SNP, consisting of increased *EEAT1* transcription, and decreased *vGLUT1* and *PSD95* transcription. Altered *vGLUT1* transcription is consistent with previous findings in mice, where Adgrl3 KO impaired vGlut1 expression^21,44^. Our data therefore support an essential role of ADGRL3 in regulating glutamatergic synapse development in humans. Interestingly, SNPs in the *EAAT1* gene have previously also been associated with ADHD, as well as several other neuropsychiatric disorders such as BPD and major depressive disorder^45^. Moreover, one specific *EAAT1* SNP associated with ADHD was found to cause increased *EAAT1* transcription^46^, consistent with the results from our study.

Lastly, scRNA-seq data generated from 8-week-old CNs was utilised to further investigate *ADGRL3* expression and genotype-associated gene expression changes. This data revealed that *ADGRL3* was primarily expressed in two cell types in our mixed cortical neuron culture; layer 2/3 neurons and cycling ventricular radial glia. This is the first time that *ADGRL3* expression has been investigated in cortical neurons at a single-cell resolution, and the data reveals further insights into its potential role in neurodevelopment. GO terms associated with *ADGRL3* expression were related to synaptic function and neuron projection, consistent with its proposed role in neurite outgrowth^12^ and synaptic development^13^. Moreover, the data revealed that rs397547 genotype altered cell type distribution, with cortical neuron cultures from risk carriers having significantly less layer 2/3 neurons and cycling ventricular radial glia. This imbalance in neuronal cell subtype could potentially lead to functional consequences such as a dysregulated excitation/inhibition balance, which has been implicated in the development of several psychiatric disorders, including ADHD^47^. Lastly, several genes previously associated with ADHD or comorbid neuropsychiatric disorders were identified as differentially expressed in layer 2/3 neurons depending on rs1397547 genotype, including *THBS1* (autism spectrum disorder^48^), *GPR37* (autism spectrum disorder^49^) and *IGF-1* (ADHD^50^).

The limitations of this study include the sample size, and that donors predominantly heterozygous for the rs1397547 SNP were used. This was largely due to the low frequency of this SNP in the general population. Future studies should aim to increase the sample size and incorporate a higher number of rs1397547 homozygous donors. Furthermore, our study was only preliminary in nature, and therefore the effects of this SNP on neuronal functionality were not investigated. Future investigations should aim to determine the functional consequences of the gene expression changes reported in this study, for example by assessing glutamatergic synapse formation and neuronal activity. Lastly, the scRNA-seq data should be interpreted with caution, as only three lines were included from rs1397547 SNP carriers, and due to overall sample size, we could not further split the groups by both diagnosis and genotype. Therefore, the results obtained may be more generalisable to an ADHD diagnosis, rather than specifically to *ADGRL3* rs1397547 genotype alone.

Taken together, our results show that the ADHD risk gene *ADGRL3* is upregulated early in human neurodevelopment and is primarily expressed in glutamatergic neurons, consistent with previous data from animal models and further supporting a role for ADGRL3 in neurogenesis and glutamatergic synapse development. We also found that the ADHD-associated SNP rs1397547 was able to significantly alter *ADGRL3* transcription at specific developmental stages. Furthermore, an altered transcriptional profile of glutamatergic synapse genes was observed in functionally mature neurons, which was specifically associated with the rs1397547 SNP independent of diagnosis. Future research should aim to further clarify the specific role of ADGRL3 in human neurodevelopment, especially as our initial data suggests that its expression is highly spatiotemporal. Moreover, the functional effects of the rs1397547 SNP on glutamatergic development should be explored, to identify potential ADHD pathomechanisms and thereby future novel drug targets.

## Acknowledgements

We thank all donors for their contribution, and Nicole Döring for the technical support. The study was supported by a NARSAD Young Investigator Grant from the BBRF, the IZKF Würzburg Clinician Scientist Program, the Open Access Publication Fund and the Graduate School of Life Sciences (both University of Würzburg). We additionally thank Dr Thomas Schubert from Nikon Germany for the microscopy support.

## Conflict of Interest

RVM, MN, FR, AC and ZS have no conflict of interest to declare. SKS has received speaker’s and author’s honoraria from Janssen, Takeda and Medice Arzneimittel Puetter GmbH&Co KG.

---

**Supplementary Figure 1: 3D localisation of ADGRL3 in another cell line.** IF labelling was performed on 2-week-old CNs and confocal microscopy performed. 3D renderings were then carried out. (A) IF labelling for TUBB3 and ADGRL3. (B) Basal distribution of ADGRL3 labelling. (C) IF labelling for vGLUT2 and ADGRL3.

**Supplementary Figure 2. ADGRL3 protein expression profile in CNs.** IF labelling was performed on 2-week-old CNs for ADGRL3, TUBB3 and vGLUT2 (n=6 CC, n=5 CG/GG). (A) The percentage of TUBB3+ regions also expressing ADGRL3 was calculated, and mean fluorescence intensity measured. (B) The same procedure was performed for vGLUT2+ regions.

**Supplementary Figure 3: Effects of rs1397547 SNP on *ADGRL3* transcription at later developmental time points.** RT-qPCR was performed to quantity relative *ADGRL3* transcription in (A) 4-week-old and (B) 12-week-old CNs.

**Supplementary Figure 4: Seurat clusters showing the different cell types present in 8-week-old CNs.** scRNA-seq was performed on 8-week-old CNs (n=9) and UMAP clustering carried out. 19 different clusters were identified using optimum parameters. Cell types were identified using publicly available brain developmental reference datasets. ccRG=cycling radial glia; ccvRG=cycling ventricular radial glia; CGE_IN=caudal ganglionic eminence interneurons; CGE_LGE_IN=caudal ganglionic eminence/lateral ganglionic eminence interneurons; INP=interneuron progenitors; IP=intermediate progenitors; L23=layer 2_3 neurons; L4=layer 4 neurons; L56=layer 5_6 neurons; L6_CThPN=layer 6 corticothalamic projection neurons; LGE-IN=lateral ganglionic eminence interneurons; mesenchyme=mesenchymal cells; oRG=outer radial glia; RG=radial glia; vRG=ventricular radial glia.

**Supplementary Figure 5. Uncropped western blot image for detecting ADGRL3 protein expression.** ADGRL3 band is visible at the estimated molecular weight of ∼139 kDa in two-week-old cortical neurons from 11 individual donors (lanes 2-12).

## Notes

### Competing Interest Statement

The authors have declared no competing interest.

