## Supplemental Information for "Expression profile of the ADHD risk gene *ADGRL3* during human neurodevelopment and the effects of genetic variation"

### Supplementary Information

#### scRNA-seq data analysis

All analyses were performed in R v. 4.2.2 using the Seurat Package. The counts matrix filtered to only include genes that were detected in at least three samples and for cells that expressed at least 200 genes. Low-quality cells with high percentage of mitochondrial (> 10%) or low ribosomal (< 5%) RNA as well as doublet cells were removed. The remaining data was then normalized (NormalizeData()), scaled (ScaleData()) and the most variable features identified (FindVariableFeatures()). The data was batch corrected and integrated using Harmony (RunHarmony()). To optimize the clustering, the data was first clustered at different resolutions (FindClusters()) and checked for cluster stability using clustree(). The most stable solution was identified at a resolution of 1.4. For visualization a UMAP with distance of 0.5 and a spread of 1 was applied. Cell cycle state inferred using CellCycleScoring().Positive (logFC>1) markers for the specific clusters were calculated using FindMarkers() and tested for GO-term enrichment using gprofiler2. Cluster Mapping to annotated cell types were done using SingleR with previously published scRNA datasets from human cerebral organoids (https://www.nature.com/articles/s41586-023-06473-y) as well as the human developing brain (https://science.org/doi/10.1126/science.adf1226). Pseudotime analyses were perfomed with the slingshot-package setting cycling radial glial cell clusters as starting point. Differentially expressed genes (DEGs) between genotypes (CC vs CG/GG) as well as between ADHD and non-ADHD were calculated using the FindMarkers() function with standard wilcoxon.test method and FDR correcting for multiple testing. DEGs were identified across all analysed cells (pseduobulk) and for each cluster, respectively. Genes with a adj.pvalue < 0.05 and a |log32FC|>0.25 were subject to gprofler2 based enrichment testing. For all enrichment tests the gene-universe was defined as the set of all genes detected in the dataset.

Cluster/Cell-type frequencies were similarly compared across diagnosis and genotype using fisher-exact testing of one Celltype vs all others; p-values were Bonferroni corrected for multiple testing.
